# Flexible and site-specific manipulation of histones in live animals

**DOI:** 10.1101/2023.03.19.533378

**Authors:** Efrat Finkin-Groner, Amni Al-Kachak, Albert Agustinus, Ryan Bastle, Ashley Lepack, Yang Lyu, Ian Maze, Yael David

**Author notes:** Equal contribution (Efrat Finkin-Groner, Amni Al-Kachak, Albert Agustinus).

## Abstract

Recent advances in protein engineering have provided a wealth of methods that allow for the site-specific manipulation of proteins *in vitro* and in cells. However, the efforts to expand these toolkits for use in live animals has been limited. Here, we report a new method for the semi-synthesis of site-specifically modified and chemically defined proteins in live animals. Importantly, we illustrate the usefulness of this methodology in the context of a challenging, chromatin bound N-terminal histone tail within rodent postmitotic neurons located in ventral striatum (Nucleus Accumbens/NAc). This approach provides the field with a precise and broadly applicable methodology for manipulating histones *in vivo*, thereby serving as a unique template towards examining chromatin phenomena that may mediate transcriptomic and physiological plasticity within mammals.

## Introduction

Chromatin regulation of gene expression is involved in numerous processes ranging from cell-cycle progression and stem cell differentiation to adulthood neural plasticity^1,2^, with abrogation of these signaling cascades often resulting in disease^3,4^. Much of the work delineating these complex phenomena has contributed to the notion of a so-called ‘histone code,’ which suggests that the combinatorial landscape of histone post-translational modifications (PTMs), along with their ‘writers’, ‘erasers’, and ‘readers’, may function as key determinants of a gene’s activity, and its potential to be activated or repressed in response to environmental stimuli^5^. Although increasing evidence has indicated the potential significance of histone PTMs in genomic regulation, we are still in the earliest stages of examining the many chromatin states involved in mammalian transcription and cell fate decisions. Furthermore, much debate still exists as to whether histone PTMs themselves are causally responsible for differences in chromatin states, and thus gene expression, or whether such differences are simply the consequence of dynamic processes associated with transcription and/or chromatin remodeling^6^. This is due to the fact that inferences of causality with respect to histone biology are often based upon correlations, as opposed to direct manipulations of the PTMs themselves.

Efforts to manipulate PTMs in histones and other proteins have been successfully performed using engineered split-inteins in a wide range of applications. For example, a tandem reaction between intein-mediated protein splicing and enzyme-mediated peptide ligation has been utilized to install PTMs as well as chemical probes onto histone H3 and other proteins including Cas9 nuclease and MeCP2 *in vitro* and *in nucleo*^7^. Moreover, split-inteins have also been utilized to introduce both PTM and photoaffinity probes to map chromatin interactome *in nucleo*^8^. Importantly, efforts to control histone PTMs has been expanded by our group and others using ultra-fast split inteins to modify both C-termini and N-termini of histones in live cells (*in cellulo*) and shown to be both robust and precise^9,10^. Despite all these milestones, no successful attempt has been reported to perform this precise histone PTM manipulation *in vivo*, which, if successful, will provide a valuable tool in the field to precisely decipher direct transcriptomics and physiological impacts of histone modifications in specific cell types and a whole organism context. Here, we report the development of a split-intein-based method to manipulate histone proteins in a site- and cell type-specific manner in mice brain and provide an essential technological leap in epigenetic research.

## Results

To develop a robust and flexible strategy for the manipulation of histones *in vivo*, we again utilize ultra-fast split inteins, which are synthetically accessible and are able to rapidly react in order to ligate polypeptides in a traceless manner. As previously mentioned, we demonstrated the utility of this approach for the generation of C-terminal modifications of histones in live cells^9^. To do so, we originally fused Int^N^ to the C-terminus of H2B and delivered a synthetic cargo fused to Int^C^, thereby producing a native-sequence, chemically-modified H2B protein in cultured cells. To further adapt this methodology for the flexible modification of histones on both unstructured termini, we began by expanding this approach to allow for *in cellulo* manipulations of the N-terminal tail of histone H3. Our design relies on the ectopic expression of Int^C^-H3, which is incorporated into chromatin by endogenous machinery, followed by the exogenous delivery of a synthetic Int^N^-carrying cargo. Since key sites of chemical modification on H3 are distributed throughout the entirety of its tail, we first generated a library of compatible pairs, in which a truncated H3 protein fused to Int^C^, is expressed, with its complementary synthetic tail fused to Int^N^ then being delivered to cells. This library allows the manipulation of residues spanning the flexible portion of the H3 tail up to residue 31 (Figure 1a). To simplify analysis of splicing reactions, we tagged Int^C^ with a Flag and Int^N^ with an HA tag. As a splicing control, we used Int^C^-fused full length H3 with endogenous exteins.

**Figure 1:**
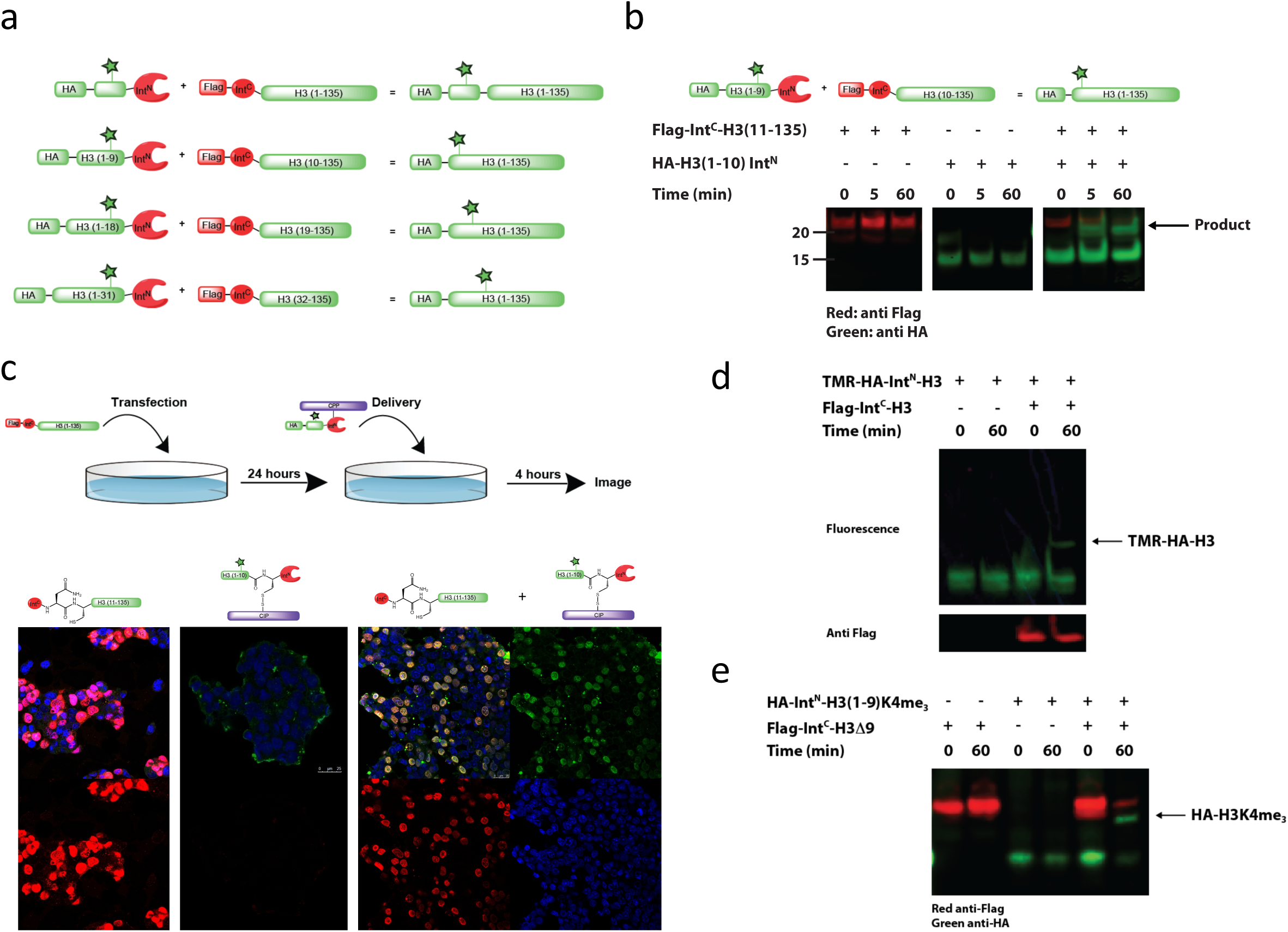
Employing protein *trans* splicing for the manipulation of histone N-terminal tails *in cellulo*. (a) Intein pairs library for the manipulation of H3 tail residues 1-31. Flag-Int^C^-H3 constructs were expressed and incorporated into chromatin, and HA-Int^N^ was delivered into cells. All splicing reactions produce a native, full-length HA-H3 with a site-specific modification (green star). (b) Representative *in nucleo* blot for chromatin associated splicing reactions. Flag-H3 (full length) was expressed in 293T cells by transfection and allowed to express for 24 hours. Cells were then harvested and hypotonically lysed to release nuclei, which were treated with 0.5 µM of HA-Int^N^ for indicated times. Samples were lysed in the presence of iodoacetamide, separated on SDS-PAGE and analyzed by western blotting with anti-Flag and anti-HA antibodies. (c) Representative IF analysis of *in cellulo* splicing reactions. Cells were transfected, as described above, and after 24 hours were treated with 2.5 µM HA-Int^N^ for 4 hours. Cells were then washed, fixed and immunostained with anti-Flag, anti-HA and DAPI. Histones can be site-specifically decorated with synthetic cargo. (d) Cells expressing Int^C^-H3 constructs were generated, as described in (b), followed by treatment with TMR-HA-Int^N^. (e) Cells expressing Int^C^-H3(Δ9) constructs were generated followed by treatment with HA-H3(1-9)K4me_3_-Int^N^. Each experiment was repeated for ≥ three times.

To test the expression and chromatin localization of the intein-fused histones, we transfected cells with each of the different constructs, followed by cellular fractionation and immunoblotting analysis. Our results indicate that all of our constructs express well and localize exclusively to chromatin, with a slight reduction in expression observed in concordance with increases in H3-tail truncation (Figure S1). To verify that the expression of intein-fused truncated H3 does not disrupt cellular functions we performed RNA-seq 24 hours post-transfection with each of the constructs tested, which revealed normal gene expression patterns in all tested samples (Figure S2).

Our design aims to minimize perturbation of ligated H3 proteins without compromising intein kinetics, which can be significantly affected by flanking extein residues^11^. To verify that the new exteins do not affect the reactivity of the new pairs on chromatin directly, we turned to *in nucleo* assays. In these assays, we hypotonically lyse transfected cells and isolate intact nuclei prior to synthetic cargo addition, which allows the uncoupling of delivery from reactivity. Since our initial Int^N^ libraries only contain a genetically encoded HA tag, we expressed and purified these proteins recombinantly achieving >95% purity (Figure S3). Each of the Int^N^ cargos was incubated with nuclei from cells expressing corresponding Int^C^ and followed the formation of full length H3 with an N-terminal HA tag, as well as loss of the Flag tagged Int^C^-H3 protein. We analyzed various time points taken from the reaction by western blotting demonstrating that all our constructs splice together to generate the same HA-H3 product with a slight decrease in efficiency of reaction observed when compared to native exteins (Figure 1b, representative blot and Figure S4, all *in nucleo* splicing reactions).

Next, to perform this reaction in live cells, we began by using directed disulfide bonding of the Int^N^ cargo to an HA2-TAT cell-penetrating peptide (CPP), which was synthesized by solid-phase peptide synthesis with an N-terminal Boc-Cys(Npys)^7^. Conjugation reactions were performed as described previously^7^ and products were purified by RP-HPLC to over 95% purity (Figure S5). This purification results in the unfolding of the Int^N^ cargo, which is a folded domain unlike Int^C^. However, previous studies have shown that Int^N^ folds back and retains its function in aqueous buffer^11^.

To test the reactivity of these conjugates, cells were first transfected with Flag-Int^C^-H3 constructs, followed by a 24-hour incubation. This allowed for the efficient incorporation of constructs throughout cell division, a major mechanism associated with histone incorporation in mitotic cells^12^. Next, 2.5 μM of Int^N^-HA cargo was delivered to cells as described before^9^. Two hours after delivery, media was changed and cells were incubated for an additional two hours (Figure 1c). Product formation, as well as chromatin localization, were evaluated using immunoflouorescence (IF) analysis. Briefly, cells were fixed, permeabilized, washed and immunoreacted with anti-Flag (starting material), anti-HA (product) and DAPI (DNA). As a control, cells were transfected with Flag-Int^C^-H3 but not delivered, while non-transfected cells were delivered with the same amount of Int^N^. Our results indicate the nuclear formation of HA-H3 product only when both intein components are present in cells (representative IF, Figure 1c, all *in cellulo* controls, Figure S6, all splicing reactions, Figure S7). It is noteworthy that disulfide bonding protects Int^N^ inteins and guarantees that the CPP is released upon endosomal escape and does not diffuse into the nucleus with its cargo.

To evaluate the utility of this methodology in splicing synthetic cargo, we synthesized a C-terminal thioester HA-peptides containing either an N-terminal organic fluorophore (tetramethylrhodamine, TMR) or an H3 tail (1-9) with K4me_3_. These peptides were ligated to recombinant N-terminal cysteine Int^N^ generated by Sumo-fusion cleaved with Ulp1 (ubiquitin like protease1). The ligation products were purified (Figure S8) and tested for reactivity *in nucleo* with their complementary transfected pairs. Results presented in Figure 1d demonstrate a similar splicing efficiency for both recombinant and synthetic cargo.

Finally, we set out to examine the possibility of effectively employing this new methodology towards the modification of chromatin-bound histones in live animals. To do so, we chose to focus our initial efforts on the ventral striatum within the central nervous system (i.e., NAc), as this brain region represents a key limbic structure of the brain’s reward circuitry, and its activity is tightly regulated through a variety of chromatin-based mechanisms, including alterations in histone PTMs, in response to environmental stimuli that, in turn, impact an animal’s behavior (e.g., exposure to drugs of abuse)^13-15^. Since neurons are post-mitotic, replication-independent, rather than -dependent, histone variants are routinely incorporated into neuronal chromatin, primarily in response to active transcription^16^. Thus, we used the replication-independent variant H3.3 to generate a Flag-Int^C^-H3.3 and Flag-Int^C^-H3.3(Δ9) constructs. We tested the expression of this new library in culture illustrating similar expression patterns in comparison to those observed with our library of canonical H3 constructs (Figure S9). We then tested the expression of the engineered Int^C^-H3.3 in NAc. To do so, we subcloned Int^C^-H3.3 constructs into a plasmid that was used for packaging into adeno-associated viral (AAV) particles (serotype 2) for neuronal specific transduction. AAV infections were used owing to their characteristically local and robust expression patterns within brain. Indeed, infecting adult mice intra-NAc with Int^C^-H3.3 resulted in its stable expression *in vivo*–colocalizing with DAPI–with its expression peaking at 21 days post-transduction, as illustrated by IF using an anti-Flag antibody (Figure S10).

Delivering synthetic peptides to animal brains is challenging in terms of penetration and stability. We thus first aimed to test whether our HA2-TAT CPP would suffice to introduce synthetic cargo into NAc neurons. We used TMR-labeled peptide, disulfide linked to HA2-TAT and directly injected 5 µl of 100 µM peptide to the ventral striatum. 30 minutes after delivery, mice were sacrificed, and brain slices were imaged for NeuN marker (neurons), TMR (cargo) and DAPI (DNA). Co-localized signal indicated that the delivered TMR cargo is internalized into neurons (see Figure S11 for representative images). Since, unlike with cultured cells, tissues cannot be washed to remove excess peptide, we next determined the half-life of Int^N^ cargos in NAc. To do so, we directly injected the same amount of peptide and assessed its presence 30-minutes *vs*. 24-hours post-delivery. IF with an anti-HA antibody indicated that non-reacting Int^N^ is internalized by 30 minutes and then cleared within 24 hours (Figure S12). This is key, as any signal that might be detected after 24 hours in the presence of both Int^N^ and Int^C^ would not reflect residual peptide, but rather would indicate the generation of stable products. To visualize product formation in live animals, we next infected mice intra-NAc unilaterally with either Flag-Int^C^-H3.3 or Flag-H3.3. 21 days later, Int^N^-HA cargo was delivered to both hemispheres (through the same cannulation site used for viral transduction), followed by an additional 24 hours of ‘incubation’ (Figure 2a). Animals were then perfused, and IF was performed for Flag, HA and DAPI. The results of this experiment, presented in Figure 2b, indicate that while Flag signal exists in both hemispheres and colocalizes with DAPI, HA signal is only observed in Int^C^-H3.3 expressing tissues indicating that H3.3 semi-synthesis can be employed in brain.

**Figure 2:**
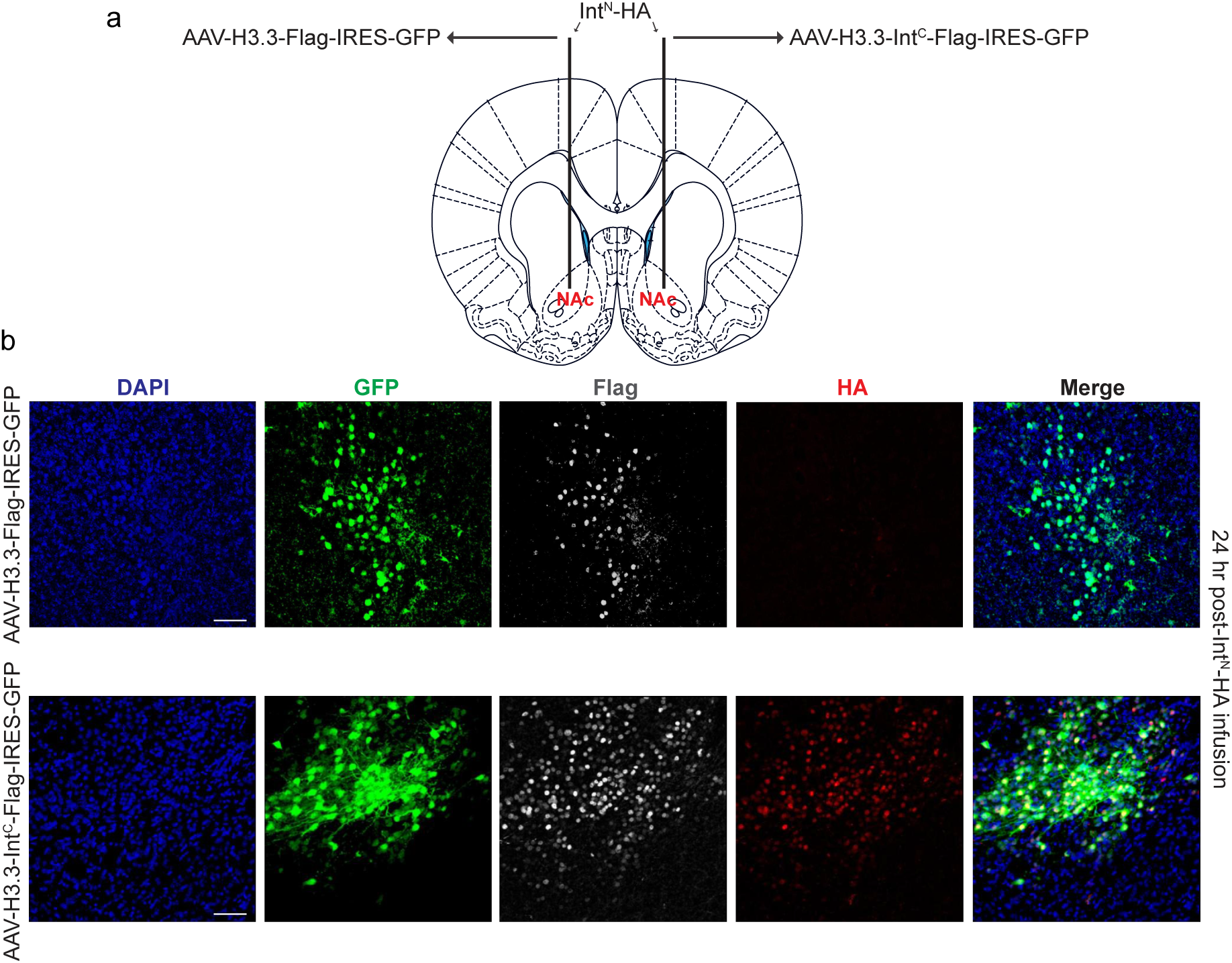
Implementation of protein *trans* splicing for manipulation of histone N-terminal tails *in vivo*. (a) Expression of Flag-Int^C^-H3.3 in mouse NAc 21 days post-AAV transduction. Mice were perfused, and brain slices were subjected to IF using an anti-Flag antibody for H3.3 incorporation, as well as DAPI for DNA and GFP for plasmid expression. (b) Product formation in NAc. Mice were injected by cannulation with AAVs expressing either Flag-H3.3 (left hemisphere) or Flag-Int^C^-H3.3 (right hemisphere). 21 days post-transduction, HA-Int^N^ was 5 µl of 100 µM peptide was injected bilaterally. 24 hours after injection, mice were perfused, and NAc slices were stained with anti-Flag, anti-HA, DAPI for DNA and GFP for plasmid expression. Each experiment was repeated for ≥ three times.

## Discussion

Altogether, we present here a flexible, robust and broadly applicable strategy for the site-specific manipulation of N-terminal histone tails. This methodology allows for the generation of chemically-defined, chromatin-bound histone proteins *in vivo*, an approach that promises to guide the field in beginning to directly assess potential causal roles for PTMs (in isolation or in combination) in regulating transcription. Using ultra-fast split inteins circumvents numerous limitations of being able to manipulate proteins at C- or N-termini, which makes them ideal for investigating functions for histone modifications *in vivo*. Since cargo are made synthetically, any solid-phase-compatible native or non-native modification can be incorporated for delivery. Moreover, using orthogonal protecting groups allows for the incorporation of multiple modifications on the same histone tail. This strategic multifunctional tool could be key to investigating “epigenetic states” established by a combination of modifications. Another advantage of this approach is that it allows for temporal control over product formation. While other genetic methodologies require days (for tissue culture) or weeks (for animal model expression), intein splicing occurs within minutes. Finally, this methodology carries with it the advantage of general applicability and robustness, as illustrated by our targeting experiments in non-replicating neurons in mice. Therefore, this approach represents an important first step in being able to resolve current conflicts in the field related to histone PTMs and their potential roles in mediating gene expression, directly or otherwise.

## Acknowledgements

The authors would like to thank Miquel Vila-Perello, as well as members of the David and Maze labs for technical and intellectual support. This work was supported by grants from the National Institutes of Health: DA044767 (I.M and Y.D.), DA042078 (I.M.), and P50 MH096890 (I.M.), as well as awards from: Josie Robertson Foundation (Y.D.), CCSG core grant P30 CA008748 (Y.D.), MQ Mental Health Research Charity, MQ15FIP100011 (I.M.) and Alfred P. Sloan Foundation Fellowship in Neuroscience (I.M.).

