## Supplementary Information for "Flexible and site-specific manipulation of histones in live animals"

[1]Chemical Biology Program, Memorial Sloan Kettering, New York, NY
[2]Department of Neuroscience, Icahn School of Medicine, Mount Sinai, New York, NY
[3]Department of Pharmacological Sciences, Icahn School of Medicine, Mount Sinai, New York, NY
[4]Department of Pharmacology, Weill Cornell Medical College, New York, NY
[5]Department of Physiology, Biophysics and Systems Biology, Weill Cornell Medical College, New York, NY
[*]Equal contribution
[§]Corresponding author

### Peptide synthesis

#### Materials

Amino acid derivatives were purchased from AGTC Bioproducts unless mentioned otherwise. Fmoc-Lys(Me3) and [5 - (and - 6) - Carboxytetramethylrhodamine] (TMR) were purchased from Anaspec. Dimethylformamide (DMF), dichloromethane (DCM), triisopropylsilane (TIS) were purchased from Fisher Scientific and used without further purification. *O*-(benzotriazol-1-yl)-*N*,*N*,*N*′,*N*′-tetramethyluronium hexafluorophosphate (HBTU), Trifluoroacetic acid (TFA), Hydrazine and *N*,*N*-diisopropylethylamine (DIPEA) were purchased from Fisher Scientific. Piperidine was purchased from Sigma Aldrich. Acetonitrile and Ether were purchased from Fisher Chemical.

Analytical reversed-phase HPLC (RP-HPLC) was performed on a Agilent 1200 series instrument with an Agilent C18 column (5 μm, 4 × 150 mm), employing 0.1% TFA in water (HPLC solvent A), and 90% acetonitrile, 0.1% TFA in water (HPLC solvent B), as the mobile phases. Analytical gradients were 0-70% HPLC buffer B over 30 min at a flow rate of 0.5 mL/min, unless stated otherwise. Preparative scale purifications were conducted on an Agilent LC system. An Agilent C18 preparative column (15-20 μm, 20 × 250 mm) or Waters semi-preparative column (12 μm, 10 mm × 250 mm) was employed at a flow rate of 20 mL/min or 4 mL/min, respectively. HPLC Electrospray ionization MS (HPLC-ESI-MS) analysis was performed on an Agilent 6120 Quadrupole LC/MS spectrometer (Agilent Technologies). UV spectrometry was performed on NanoDrop 2000c (Thermo Scientific).

#### General method

Standard Fmoc-based Solid Phase Peptide Synthesis (Fmoc-SPPS) was used for the synthesis of all peptides in this study. All peptides were synthesized on ChemMatrix resins with either 2-Chlorotrityl Chloride (CTC) or rink amide linkers (Biotage). The CTC resin was first dehydrated with DMF for 20min followed by two 1h activation with hydrazine/DIPEA four- and eight-fold excess respectively. Peptides were synthesized using manual addition of the reagents (using a stream of dry N2 to agitate the reaction mixture). For amino acid coupling, 5 eq. Fmoc protected amino acid were pre-activated with 4.9 eq. HBTU, 4.9 eq. HOBt and 10 eq. DIPEA in DMF and then reacted with the C-terminally deprotected peptidyl resin. The resin was washed with DMF multiple times, and reaction completion was monitored using the Kaiser test (Anaspec). Fmoc deprotection was performed in 20 % (v/v) piperidine in DMF, and the deprotected peptidyl resin was washed thoroughly with DMF to remove trace piperidine. Double coupling was performed for all Pro, Arg and Lys addition. Cleavage from the resin and side-chain final deprotection were performed with 95 % TFA, 2.5 % TIS and 2.5 % H2O at room temperature for 2 hours. Peptide were then precipitated with cold diethyl ether, isolated by centrifugation, dissolved in water with 0.1 % TFA and analyzed via RP-HPLC and ESI-MS. Primary sequences of synthetic peptides prepared are shown in Table S1.

##### Synthesis of C-terminal thioester peptides

HA-H3(1-9)K4Me3 synthesis was done as described above. Double coupling was performed for all trimethyl lysine.

TMR peptide synthesis

TMR coupling to the N terminus was done overnight protected from light as described above. Cleavage from the resin and side-chain deprotection were performed with 95 % TFA, 2.5 % TIS and 2.5 % $H_2O$ at room temperature for 2 hours.

Synthesis of HA2-TAT

All $Ava^N$ peptides were delivered into cultured cells by conjugation to a cell penetrating peptide (CPP) derived from the viral protein transduction domains HA2 and TAT (PMID: 17923117).

The HA2-TAT was synthesize as described before (PMID: 25901817). Briefly, standard Fmoc-based SPPS was used with Rink Amide resin (Biotage) to generate C-terminal amides. Boc-Cys(Npys)-OH (Bachem) was coupled to the N-terminus to generate active cysteine. The final peptide was purified and analyzed by RP-LC-ESI-MS.

### Histone modification in culture

#### Materials

All reagents were purchased from TFisher-Thermo Scientific unless otherwise noted. For cloning, T4 Polynucleotide Kinase (T4 PNK), Phusion High-Fidelity DNA Polymerase kit, T4 DNA ligase kit, and DPNI were used. Restriction enzymes were obtained from New England BioLabs. Primer synthesis and DNA sequencing were performed by Integrated DNA Technologies and Genewiz, respectively. PCR amplifications were performed on a Bio-Rad T100™ Thermal Cycler. PCR purification columns were purchased from Qiagen. EDTA and cOmplete, Mini Protease Inhibitor Cocktail were purchased from Sigma Aldrich. Profinity™ IMAC Resin, Ni-NTA beads were purchased from BIORAD. Centrifugal filtration units were purchased from Sartorius, and MINI dialysis units purchased from Pierce.

#### Cloning

The Flag-$Int^C$-H3.1 sequence (Flag-$Npu^C$-H3.1) was cloned into the pEGFP-N1 vector. In order to obtain the option of inserting PTM at various positions on the H3 tail, four different truncations were cloned (2.2-2.5 table S2) using primers listed in table S3.

For cargo delivery, HA-$Int^N$-His (HA-$Ava^N$-His) was cloned into a pET3 vector for bacterial expression (PMID: 25901817). Four different truncations complementary to Flag-$Npu^C$-H3.1 constructs were cloned (Table S4). Constructs S4.2 and S4.5 were obtain using HA-$Ava^N$-His as template.

#### Purification of HA-H3(1-x)-$Ava^N$-His

BL-21 cells were transformed with the expression plasmids grown at 37 °C under ampicillin selection in LB media to OD600 = 0.6 and then induced with 1mM final IPTG at 37 °C for 12 hr. After pelleting cells were resuspended in 20 mL of Lysis buffer (25 mM HEPES, 150 mM NaCl, and 1 mM DTT), supplemented with protease inhibitors and lysed by sonication. Lysates were spun down and His-tagged proteins were extracted from the insoluble fraction using extraction buffer (6 M guanidine, 20 mM Tris Ph 7.5, 1 mM EDTA and 1 mM DTT). Purification was performed using Ni-NTA beads with proteins eluted with 5 mL of elution buffer (6 M guanidine, 20 mM Tris, 1 mM EDTA, 1 mM DTT and 0.5 M imidazole) and dialyzed against 6 M guanidine, 20 mM Tris, 1 mM EDTA, 1 mM DTT. For *in nucleo* experiments, proteins were concentrated and refoldad step-wise (4 M and then 2 M guanidine buffer). The purified constructs were analyzed and quantified by RP-LC-ESI-MS.

**Purification of His-SUMO-Cys-Ava$^N$**

Construct was expressed in BL-21 as described above. Cells were pelleted and resuspended in 15 mL of extraction buffer (6 M guanidine, 20 mM Tris Ph 7.5, 1 mM EDTA and 1 mM DTT) and purified on Ni-NTA beads as described above. Protein was refolded by step-wise dialysis using 100 mM Arginine 20 mM Tris pH 7.5, 1 mM EDTA, 2 mM DTT with 4 M and then 1.5 M urea. SUMO was cleaved by the addition of 100 µg/ml Ubl-specific protease 1 (ULP1) overnight at 4$^o$C. Next, solid urea was added to 6 M and cleaved cys-Ava$^N$ was recovered by reverse Ni-NTA purification (collecting the unbound fraction). Finally, the protein was purified by RP-HPLC and fractions were analyzed by RP-LC-ESI-MS. Pure fractions were pooled, aliquoted and lyophilized.

**Ligating peptide thioester cargo to Ava$^N$**

The final products TMR-HA-AVA$^N$ and HA-H3(1-9)K4Me3-AVA$^N$ were generated by expressed protein ligation (EPL) using cys-AVA$^N$ and the corresponding synthetic peptides described in table S1.

EPL was initiated by dissolving 600 nmoles of peptide in ligation buffer (6 M guanidine, 300 mM phosphate buffer, degassed with N2 and adjust to pH 3). 10 mM of Sodium nitrate ($NaNO_2$) was added to the reaction mixture and incubated on ice for 10 min. MPAA buffer (33.6g/L MPAA in 6 M guanidine) was added to the reaction mixture and incubated for 45 min at room temperature with gentle mixing. Next, 50 mmoles of cys-Ava$^N$ were dissolved in 60 µl TCEP buffer (100 mM TCEP, 600 mM guanidine, pH 7), added to the peptide solution and pH was adjusted to 7. The reaction was allowed to proceed overnight at room temperature and the ligated product was isolated by RP-HPLC purification using a 25-60% solvent B gradient. Pure fractions were pooled, aliquoted and lyophilized. The purified products were analyzed by RP-LC-ESI-MS (see Table S2).

**Conjugation of HA2-TAT to Ava$^N$**

HA2-TAT conjugation was performed as described before (PMID: 25901817). In brief, 50 nmoles of each final EPL product were resuspend in 100 µl of 6 M guanidine buffer, 20 mM Tris, PH 7.5, 1 mM DTT. Samples were sonicated for 10 min and left at room temperature for 1 hour to reduce any spontaneous disulfide bridges. Next, reaction was diluted with 500 µl reaction buffer (6 M guanidine, 100 mM phosphate buffer, pH 6), sonicated and added to 200 nmoles of HA2-TAT. Reaction was sonicated and incubated at room temperature for 1 hour. This step was repeated twice with an additional batch of 200 nmole HA2-TAT. The final conjugated product was isolated by RP-HPLC purification using a 45-75% solvent B gradient. Pure fractions were pooled, aliquoted and lyophilized. The purified peptides were analyzed by RP-LC-ESI-MS (Table S2).

### Transfections

#### Materials

All cell culture, transfection reagents and chemicals were obtained from Thermo Fisher Scientific unless otherwise indicated. HyClone™ Dulbecco's Modified Eagles Medium High glucose without L-glutamine, sodium pyruvate (DMEM) was obtained from GE Healthcare. Fetal bovine serum (FBS) was purchased from memorial Sloan Kettering core facilities. Iodoacetamide and L-Glutamine were purchased from Acros Organics.

#### Transfection

293T cells (ATCC) were cultured in DMEM supplemented with 10 % FBS, 2 mM L-Glutamine, and 500 units/ml penicillin and streptomycin. Transfections were performed using lipofectamine 2000 following manufacture instructions. Transfected 293T cells were harvested on ice in cold PBS and kept in liquid nitrogen until use.

### Western blotting

All western blots were performed using PVDF membranes (Bio Rad), primary antibodies annotated in Table 4 and Li-COR secondary antibodies annotated in Table 5 following manufacturer instructions. Blots were imaged on Odyssey CLx Imaging System.

### *In nucleo* assay

Nuclei was isolated as previously described (PMID: 25901817). $Int^N$ constructs were incubated with 90 mM TCEP for 30min prior to reaction to reduce any spontaneously formed disulfide bonds and then added to nuclei at the designated concentrations (0.05-1 μM). Reactions were allowed to proceed for the indicated times at 37 °C before being quenched by the addition of iodoacetamide to a final concentration of 80 mM. For SDS-PAGE analysis, the reaction mixtures were sonicated twice for 25 seconds with rod sonicator (Fisher Scientific) and boiled for 10 minutes with SDS sample buffer. Each experiment was repeated for ≥ three times.

### Peptide delivery and immunofluorescent (IF) staining

Approximately 230,000 293T cells were plated on a 12 well glass coverslip and grown for 24 hours. Cells were then transfected with Flag-$Npu^C$-H3 constructs as described above. After additional 24 hours, cells were washed with OPTI-MEM minimal media and incubated with 5 μM of $Int^N$ cargo for 2 hours. After two hours and equal volume of full media was added, and cells were incubated for additional 2-3 hours before proceeding to IF staining.

Cells stained by IF using 100% Methanol for fixation, 0.2% triton x-100 for permeabilization and 1% BSA, 10% goat serum, 0.02% NaAz in PBS for blocking. Primary and secondary antibodies were used in the dilution indicated in Table 4 and 5. DAPI stain (Thermo Fisher Scientific) was used according to manufacturer protocol. Cell were imaged under confocal microscopy at 63x magnification using a Leica SP5 microscope (Leica Microsystems GmbH, Germany). Several cross-section images were taken per experimental condition. Each experiment was repeated for ≥ three times.

### Histone modification in mice

#### Animals

Male C57BL/6J mice (~25 g) were purchased at 8 weeks of age from the Jackson Laboratory and allowed 1 week of acclimation to the Icahn School of Medicine at Mount Sinai (ISMMS) housing facilities before the start of experiments. All mice were group-housed and maintained on a 12h light/dark cycle with food and water available ad libitum. All mouse procedures were performed in accordance with the National Institutes of Health Guide for Care and Use of Laboratory Animals and the ISMMS Animal Care and Use Committee.

#### Stereotaxic surgeries

Mice were anesthetized with a mixture of ketamine (100 mg/kg) and xylazine (10 mg/kg) and positioned in a small-animal stereotaxic instrument (Kopf Instruments), and the skull surface was exposed. 26s gauge syringe needles (Hamilton) were used to bilaterally infuse 0.5 µl of virus/peptide into the nucleus accumbens (NAc) at a rate of 0.1 mL/min.

##### CPP infusion experiment

CPP fused with TMR was generated as 1 µM peptide in 100% DMSO and diluted to 100 µM peptide in 10% DMSO and injected into NAc at a rate of 0.1 µL/min for 5 minutes (bregma coordinates: anterior/posterior 1.6 mm; medial/lateral, 1.5mm; dorsal/ventral. -4.4 mm; 10° angle). Animals were perfused 30 minutes after viral infusion to confirm peptide penetration *in vivo*.

##### Peptide washout experiment

HA-H3 (1-10)-IntN was generated as 5 mM peptide in 100% DMSO and diluted to 100 µM peptide in 2% DMSO and injected into NAc at a rate of 0.1 µL/min for 5 minutes (bregma coordinates: anterior/posterior 1.6 mm; medial/lateral, 1.0mm; dorsal/ventral. -4.5 mm). Animals were perfused 30 minutes after infusion to confirm peptide penetration in vivo and 24 hours after infusion to confirm peptide washout.

##### Intein reaction experiment

22 gauge bilateral guide cannulae (Plastics One, #305176) were implanted into NAc (bregma coordinates: anterior/posterior 1.6 mm; medial/lateral, 1.5mm; dorsal/ventral. -4.4 mm). AAV expressing H3.3-IntC-Flag-IRES-GFP and AAV-H3.3-Flag-IRES-GFP was infused at a rate of 0.1

μL/min for 5 minutes using a Harvard Apparatus PicoPlus Syringe Pump (70-2213). To allow for optimal AAV expression, peptides were infused three weeks post-viral infusion. HA-H3 (1-10)-IntN was generated as 5 mM peptide in 100% DMSO and diluted to 100 μM peptide in 2% DMSO and infused at a rate of 0.1 μL/min for 5 minutes using a Harvard Apparatus PicoPlus Syringe Pump. Animals were perfused 24 hours after peptide infusion. Each experiment was repeated for ≥ three times.

### Immunohistochemistry

Mice were anesthetized with a mixture of ketamine (100 mg/kg of body weight) and xylazine (10 mg/kg of body weight) and perfused with 1x PBS and 4% paraformaldehyde in 1x PBS at a pH of 7.4. Intact brains were removed and stored in 4% PFA for 24h and transferred to PBS containing 30% sucrose for 72h before being sectioned at 40 μm.

<u>CPP infusion experiment</u>

Free floating NAc sections were washed with 1X PBS and then incubated with 1X PBS containing DAPI (1:10,000) Sections were once again washed, followed by mounting with ProLong Gold Antifade. All sections were imaged using standard fluorescence or confocal microscopy.

<u>Peptide washout experiment</u>

Free floating NAc sections were washed with 1X PBS, blocked (3% normal goat serum, 0.1% TritonX-100, 1X PBS) and later incubated with anti-HA antibody (Table 4) in blocking solution. Following overnight incubation, sections were rinsed three times for 10 minutes with 1X PBS, followed by incubation with Alexa Fluor 488 donkey anti-rabbit secondary antibodies in 1X PBS blocking solution for two hours (Table 5). Nuclear co-staining was achieved by incubating sections in 1X PBS containing DAPI (1:10,000) for 10 minutes. Sections were once again washed, followed by mounting with ProLong Gold Antifade. All sections were imaged using standard fluorescence or confocal microscopy.

<u>Intein reaction experiment</u>

Free floating NAc sections were washed with 1X PBS, blocked (3% normal goat serum, 0.1% TritonX-100, 1X PBS) and later incubated with anti-HA antibody, anti-Flag antibody, and anti-GFP antibody in blocking solution (Table 4). Following overnight incubation, sections were rinsed three times for 10 minutes with 1X PBS, followed by incubation with Alexa Fluor 568 donkey anti-rabbit, Alexa Fluor 647 donkey anti-mouse, and Alexa Fluor 488 donkey anti-rabbit secondary antibodies in 1X PBS blocking solution for two hours (Table 5). Nuclear co-staining was achieved by incubating sections in 1X PBS containing DAPI (1:10,000) for 10 minutes. Sections were once again washed, followed by mounting with ProLong Gold Antifade. All sections were imaged using standard fluorescence or confocal microscopy.

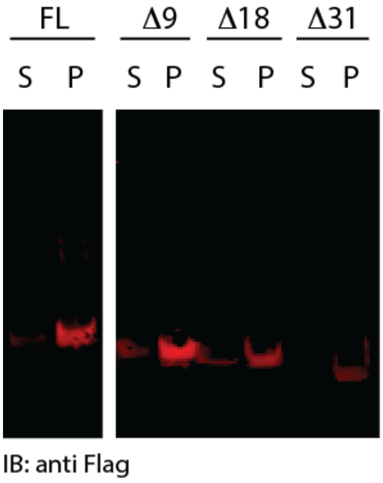


**<u>Figure S1:</u> Expression and chromatin localization of all the Flag-Int$^{C}$-H3ΔX constructs**. Constructs were transfected and expressed under a constitutive CMV promoter. 24 hours after transfection cell were harvested and fractionated to soluble and chromatin fraction and blotted with anti-Flag. Blots showing chromatin-localization and a slight decrease in expression levels with increased H3 truncation.

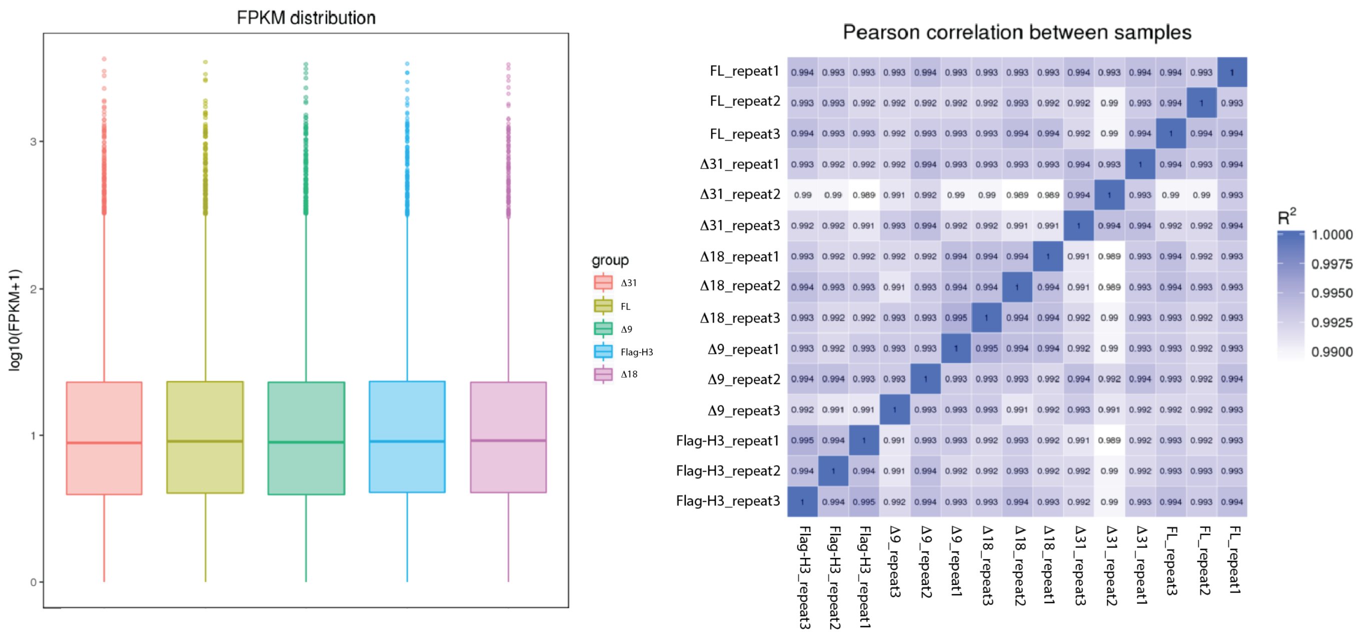


<u>**Figure S2:**</u> **RNA-seq of all the Flag- H3 and Flag-H3ΔX constructs**. Constructs were transfected and expressed under a constitutive CMV promoter. 24 hours after transfection cell were harvested, RNA was extracted and sequenced. RNA-seq results indicate perturbation is equivalent to the expression of a flag tagged H3.

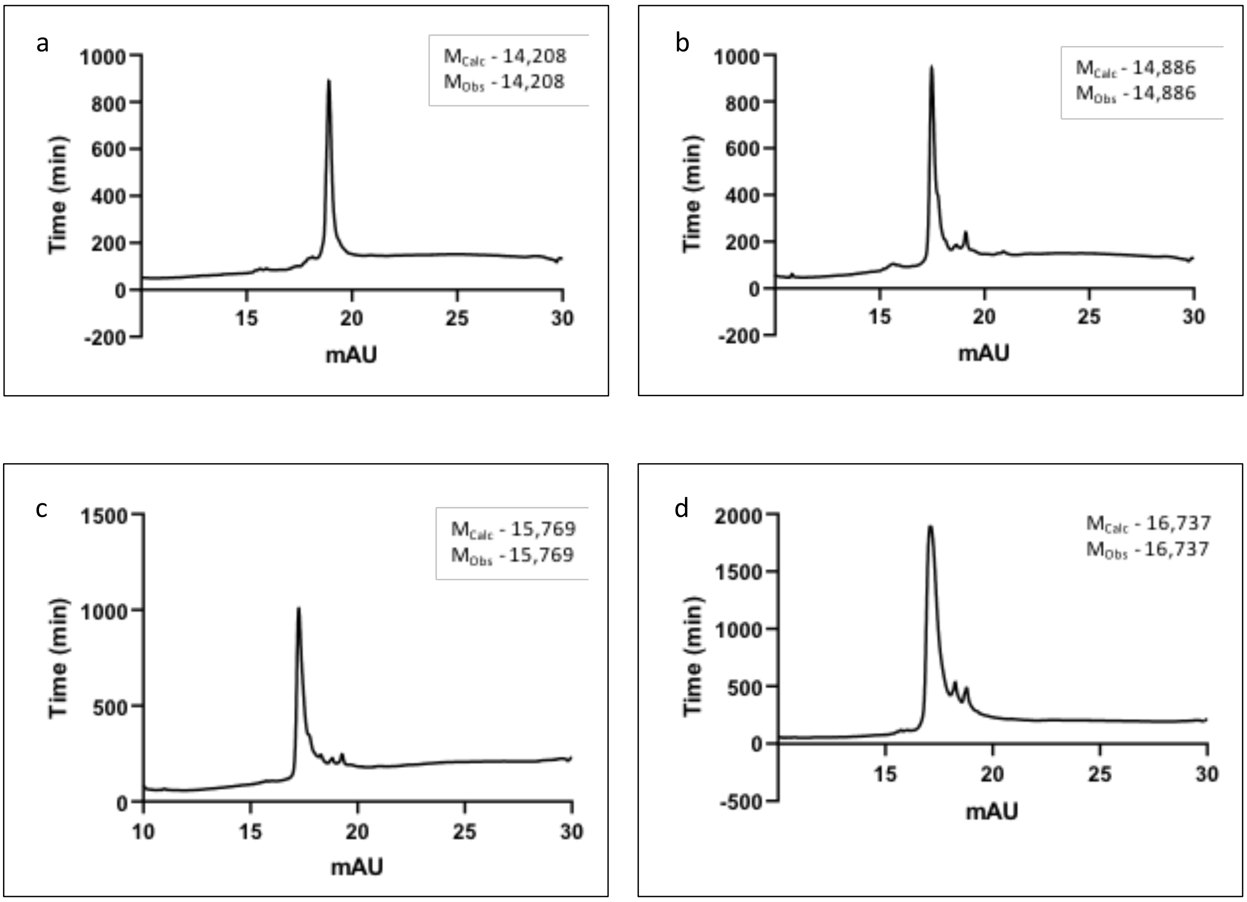


**<u>Figure S3:</u> Purification of HA-Int$^{N}$-H3 (1-X) constructs**. Constructs were transformed and expressed in E.coli as described above. Purity was evaluated by RP-LC/MS with final products over 95% purity used in the assays. Estimated and observed mass indicated for each construct. (a) HA-Int$^{N}$, (b) HA-H3(1-9)-Int$^{N}$, (c) HA-H3(1-18)-Int$^{N}$, (d) HA-H3(1-31)-Int$^{N}$.

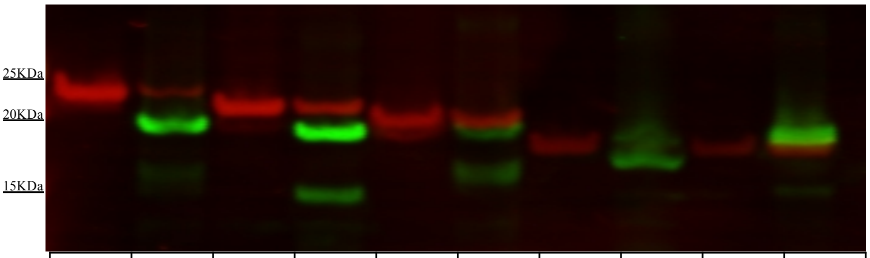


| | | | | | | | | | |
|---|---|---|---|---|---|---|---|---|---|
| **0** | **60** | **0** | **60** | **0** | **60** | **0** | **60** | **0** | **60** |
| FL-$Int^{N}$H3<br>+FL-$Int^{C}$ H3 | | $Int^{N}$ H3(1-9)<br>+$Int^{C}$H3($\Delta$9) | | $Int^{N}$ H3(1-18)<br>+$Int^{C}$ H3($\Delta$18) | | $Int^{N}$H3(1-27)<br>+$Int^{C}$ H3($\Delta$27) | | $Int^{N}$ H3(1-31)<br>+$Int^{C}$H3($\Delta$31) | |

**<u>Figure S4:</u> *In nucleo* semi-synthesis of HA-H3**. Indicated Flag-$Int^{C}$ constructs were transfected and 24 hours later harvested. Nuclei were isolated and incubated with 0.5 µM of the corresponding HA-$Int^{N}$ constructs for 60 minutes. Samples were then quenched and analyzed by western blot with anti-Flag (red) and anti-HA (green).

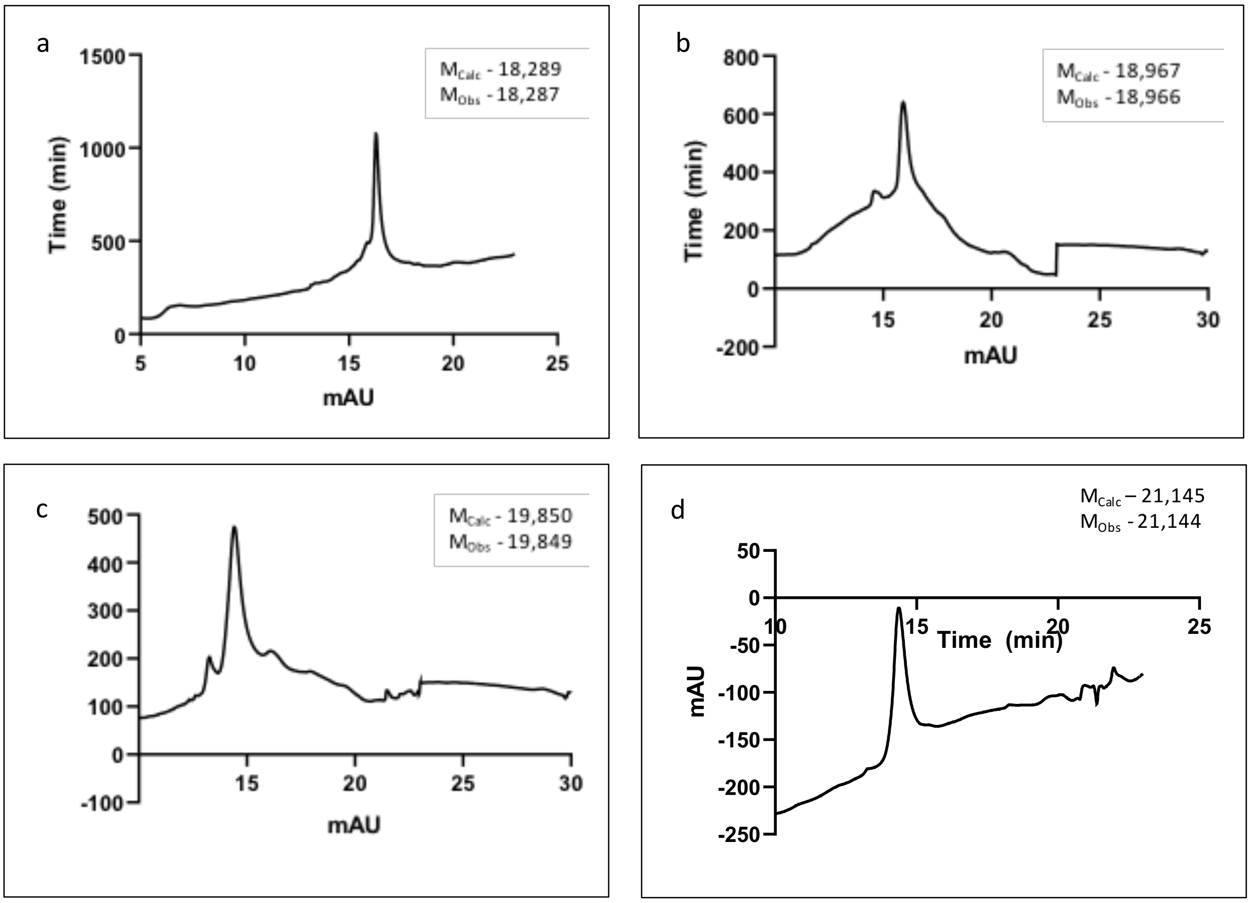


**<u>Figure S5:</u> Purification of HA-Int$^{N}$-H3 (1-X)~HA2-TAT constructs**. Int$^{N}$ constructs were purified as described above. HA2-TAT cell penetrating peptides was synthesized using solid-phase peptide synthesis. Directed disulfide conjugation was performed as described and was followed by RP-HPLC purification. Pure fractions were pooled and evaluated by RP-LC/MS with final products over 95% purity used in the assays. Estimated and observed mass indicated for each construct. (a) HA-Int$^{N}$~HA2-TAT, (b) HA-H3(1-9)-Int$^{N}$~HA2-TAT, (c) HA-H3(1-18)-Int$^{N}$~HA2-TAT, (d) HA-H3(1-31)-Int$^{N}$~HA2-TAT.

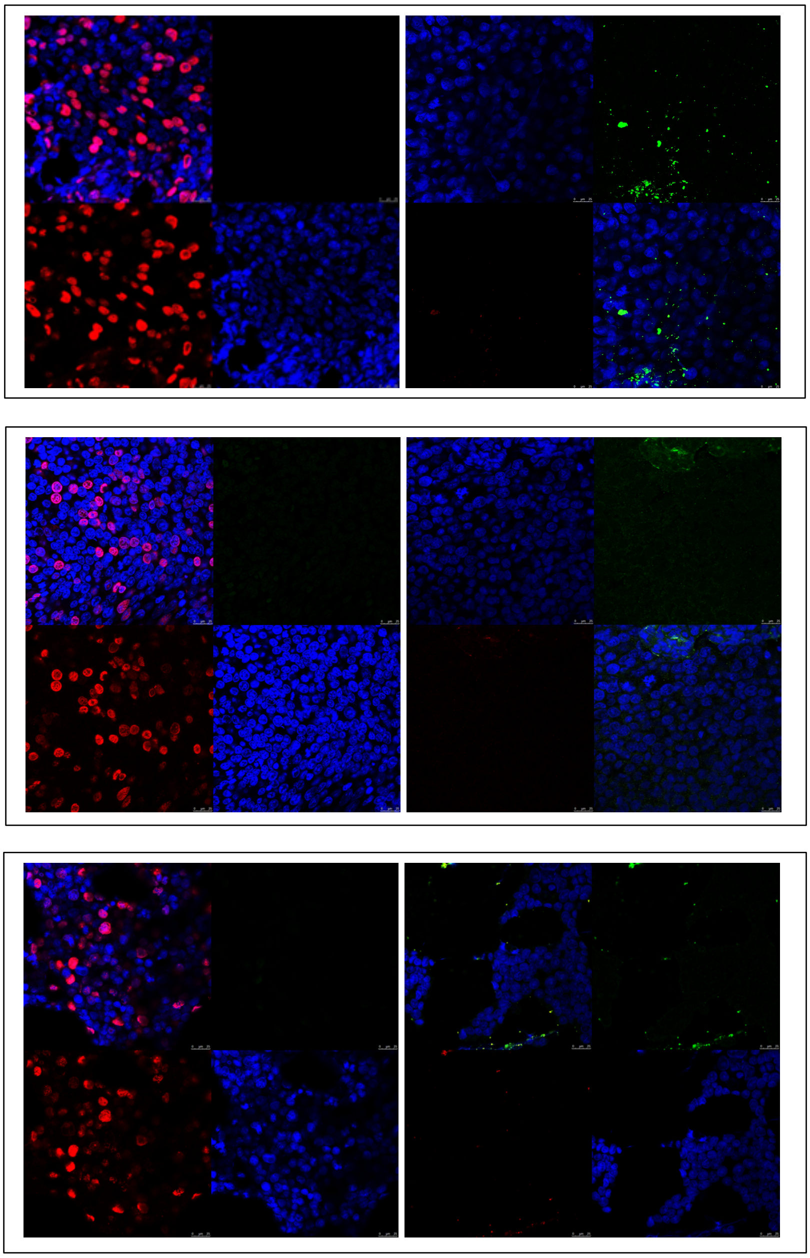

**<u>Figure S6:</u> Representative immunofluorescence images of control experiments for in vivo splicing experiments**. Cell were either transfected (left) or delivered (right) as described above. Immunofluorescence was performed using anti-Flag and anti-HA to both conditions for each of the following pairs: Flag-Int$^{C}$-H3$\Delta$9 and HA-Int$^{N}$-H3(1-9)~HA2-TAT (top), Flag-Int$^{C}$-H3$\Delta$18 and HA-Int$^{N}$-H3(1-18)~HA2-TAT (middle), Flag-Int$^{C}$-H3$\Delta$31 and HA-Int$^{N}$-H3(1-31)~HA2-TAT (top)

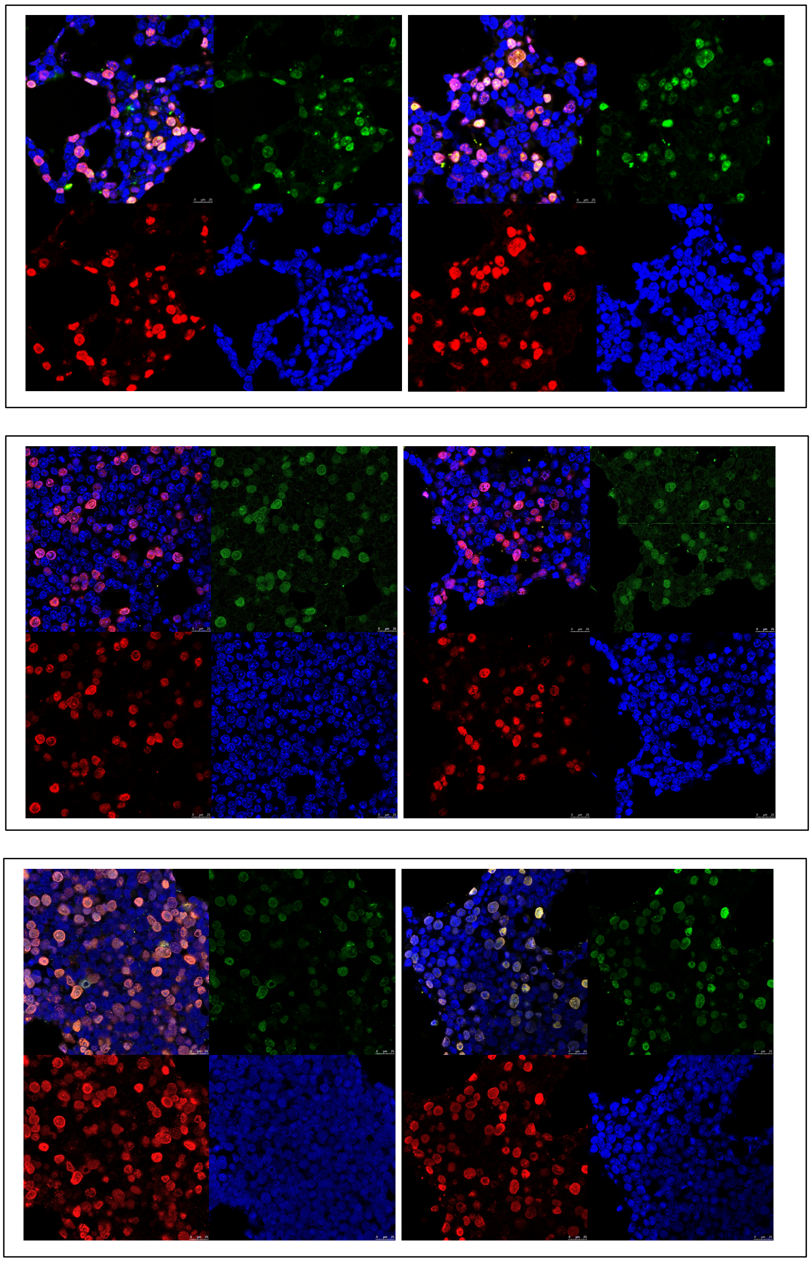

**<u>Figure S7:</u> Representative immunofluorescence images of cells transfected and delivered with the compatible intein pairs**. Cell were transfected and delivered as described above. Two representative experiments are presented for each of the following pairs: Flag-Int$^{C}$-H3Δ9 + HA-Int$^{N}$-H3(1-9)~HA2-TAT (top), Flag-Int$^{C}$-H3Δ18 + HA-Int$^{N}$-H3(1-18)~HA2-TAT (middle), Flag-Int$^{C}$-H3Δ31 + HA-Int$^{N}$-H3(1-31)~HA2-TAT (top)

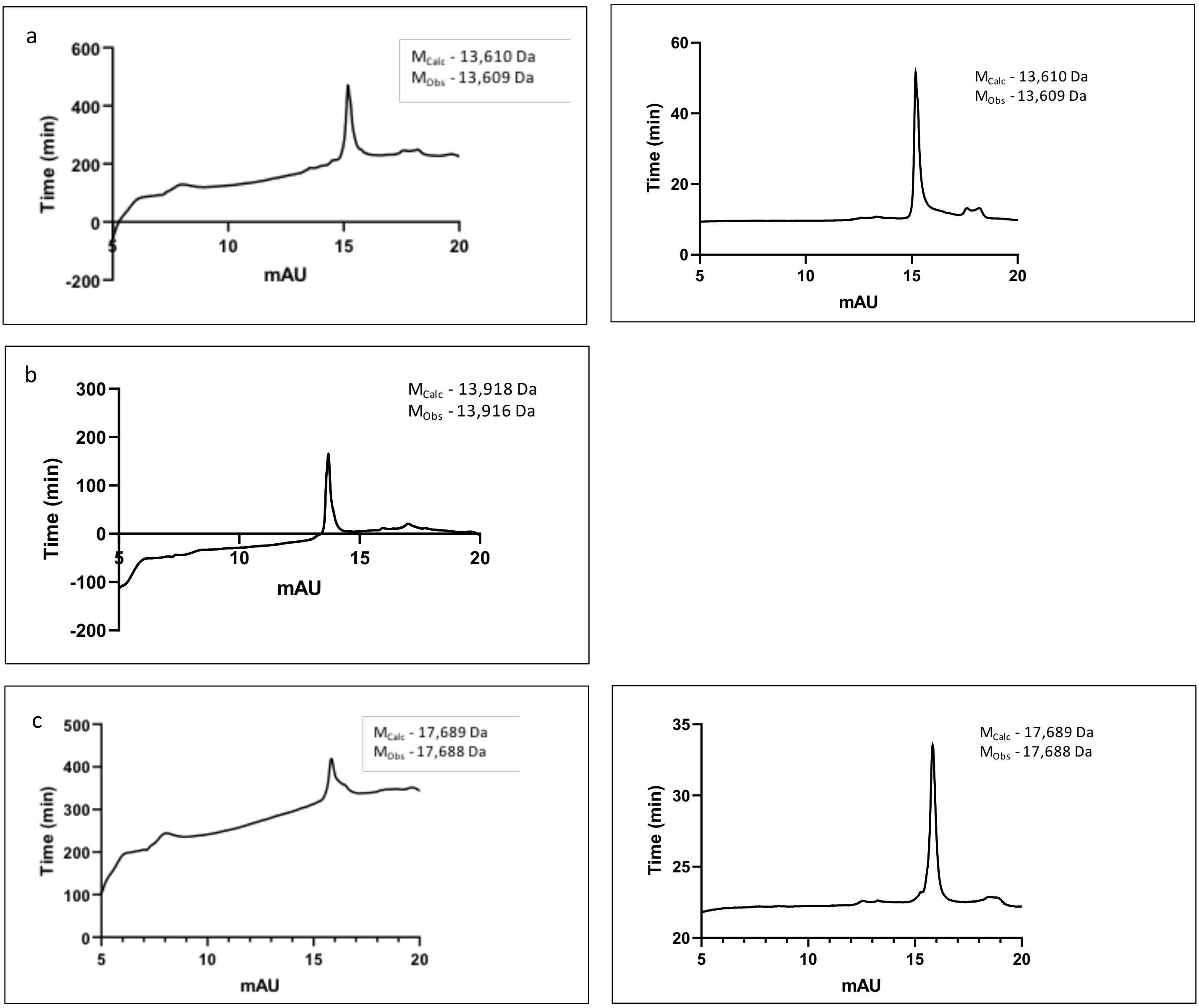


**<u>Figure S8:</u> Generated semi-synthesis HA-Int$^{N}$ constructs.** Peptides were ligated to Int$^{N}$ and purified as described above. Pure fractions were pooled and evaluated by RP-LC/MS with final products over 95% purity used in the assays. Estimated and observed mass indicated for each construct. (a) TMR-HA-Int$^{N}$ analyzed using 214 (left) and 540 (right) wavelengths (b) HA-H3(1-9)K4me3-Int$^{N}$ (c) TMR-HA-Int$^{N}$~HA2-TAT analyzed using 214 (left) and 540 (right) wavelengths.

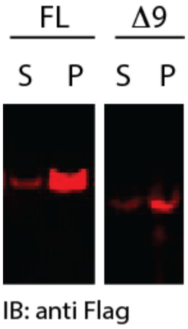


**<u>Figure S9:</u> Expression and chromatin localization of Flag-Int$^{C}$-H3.3 constructs**. Constructs were transfected and expressed under a constitutive CMV promoter. 24 hours after transfection cell were harvested and fractionated to soluble and chromatin fraction and blotted with anti-Flag. Blots showing chromatin-localization and a decrease in expression levels with increased H3 truncation.

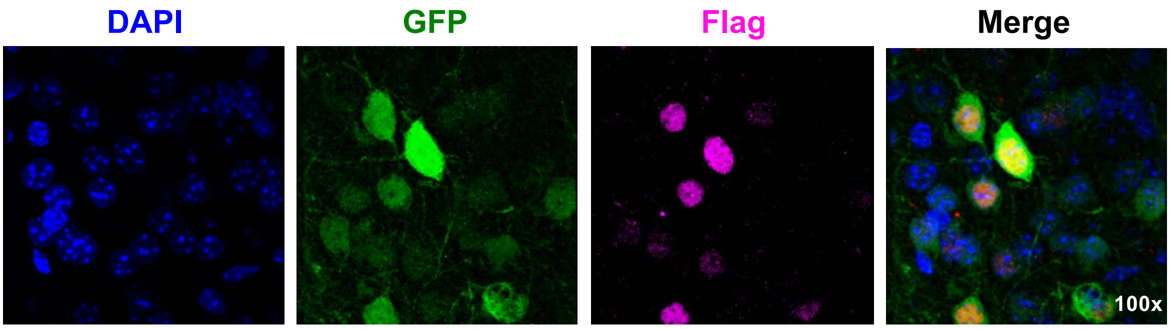


AAV-Flag-Int$^{C}$-H3.3 (1-135)-IRES-GFP
delivered intra-Nac

**<u>Figure S10:</u> Histone variant-intein fusions can be efficiently expressed and incorporated into neuronal chromatin in adult brain.** Lentiviral mediated expression of an H3.3-Int$^{N}$-Flag vector in adult mouse NAc. Brain sections were co-stained with DAPI (nuclear), GFP (viral expression) and Flag (intein-fused H3.3).

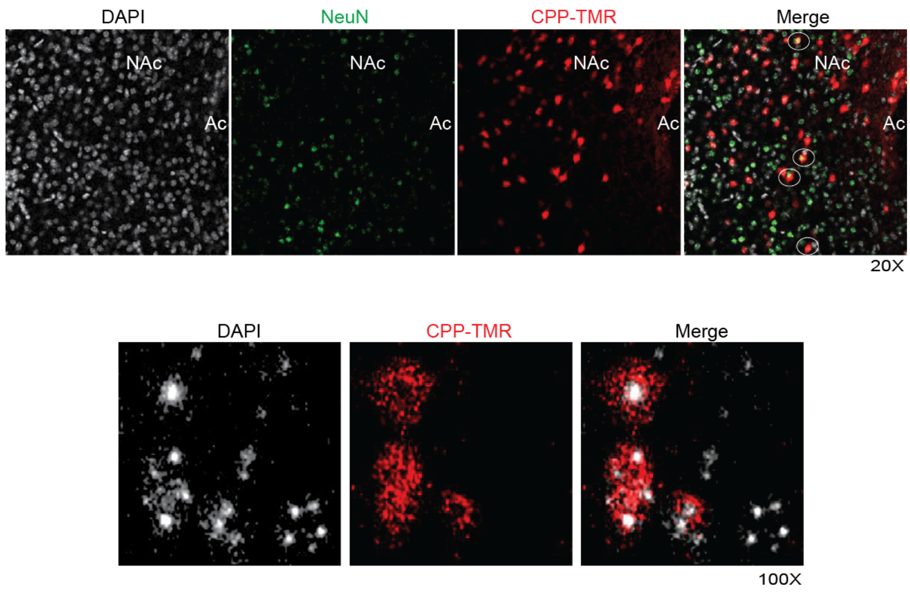


**<u>Figure S11:</u> Delivery of synthetic CPP-TMR into adult mouse NAc.** Peptide was delivered by direct injection showing penetration and nuclear localization of the synthetic peptide. Brain sections were imaged for TMR and co-stained with DAPI (nuclear) and NeuN (neuronal marker).

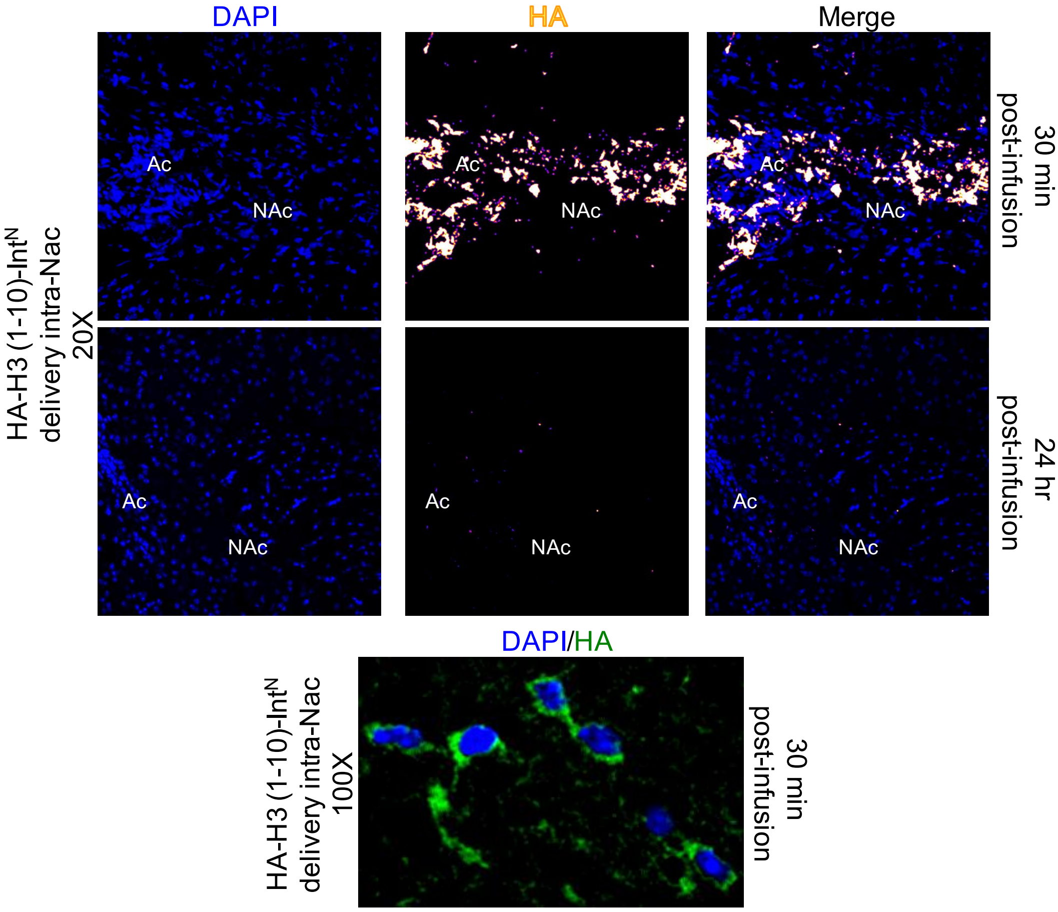


**<u>Figure S12:</u> Unreacted peptide is cleared within 24 hours.** Peptide was delivered as described above and brain sections were co-stained with DAPI (nuclear) and anti-HA (peptide) after 30 minutes and 24 hours.

Table S1 - Sequences of synthesized peptides

| **Synthetic peptide** | **Amino acid sequence** | **Observed mass (Da)** | **Expected mass (Da)** |
|---|---|---|---|
| TMR-HA | GYPYDVPDYAGGAEY | 2062 | 2064 |
| HA-H3(1-9) K4Me3 | GYPYDVPDYAGGARTK(Me)3QTARK | 2370 | 2371 |
| Cys(Npys)-HA2-TAT | Cys(Npys)-CGLFEAIAEFIENGWEGLIEGWYGGRKKRRQRRR | 4235 | 4235 |

Table S2 - Amino acid sequences of the various truncations of intein constructs

| **Gene** | **Amino acid sequences** | **Observed mass (Da)** | **Expected mass (Da)** |
|---|---|---|---|
| Flag-Npu$^C$-H3(1-135) | MGDYKDDDDKGIKIATRKYLGKQNVYDIGVERDHNFALKNGFIASNCFNARTKQTARKSTGGKAPRKQLATKAARKSAPATGGVKKPHRYRPGTVALREIRRYQKSTELLIRKLPFQRLVREIAQDFKTDLRFQSSAVMALQEACEAYLVGLFEDTNLCAIHAKRVTIMPKDIQLARRIRGERA | - | - |
| Flag-Npu$^C$-H3(Δ9) | MGDYKDDDDKGIKIATRKYLGKQNVYDIGVERDHNFALKNGFIASNCFGGKAPRKQLATKAARKSAPATGGVKKPHRYRPGTVALREIRRYQKSTELLIRKLPFQRLVREIAQDFKTDLRFQSSAVMALQEACEAYLVGLFEDTNLCAIHAKRVTIMPKDIQLARRIRGERA | - | - |
| Flag-Npu$^C$-H3(Δ18) | MGDYKDDDDKGIKIATRKYLGKQNVYDIGVERDHNFALKNGFIASNCLATKAARKSAPATGGVKKPHRYRPGTVALREIRRYQKSTELLIRKLPFQRLVREIAQDFKTDLRFQSSAVMALQEACEAYLVGLFEDTNLCAIHAKRVTIMPKDIQLARRIRGERA | - | - |
| Flag-Npu$^C$-H3(Δ28) | MGDYKDDDDKGIKIATRKYLGKQNVYDIGVERDHNFALKNGFIASNCFAATGGVKKPHRYRPGTVALREIRRYQKSTELLIRKLPFQRLVREIAQDFKTDLRFQSSAVMALQEACEAYLVGLFEDTNLCAIHAKRVTIMPKDIQLARRIRGERA | - | - |
| Flag-Npu$^C$-H3(Δ31) | MGDYKDDDDKGIKIATRKYLGKQNVYDIGVERDHNFALKNGFIASNCVGVKKPHRYRPGTVALREIRRYQKSTELLIRKLPFQRLVREIAQDFKTDLRFQSSAVMALQEACEAYLVGLFEDTNLCAIHAKRVTIMPKDIQLARRIRGERA | - | - |
| HA-AVA$^N$ | MYPYDVPDYAGGAEYCLSYDTEVLTVEYGFVPIGEIVDKGIESSVFSIDSNGIVYTQPIAQWHHRGKQEVFEYSLEDGSIIKATKDHKFMTQDGKMLPIDEIFEQELDLLQVKGLPEGGHHHHHH | 14208 | 14208 |
| HA-H3(1-9)-AVA$^N$ | MYPYDVPDYAGGARTKQTARKCLSYDTEVLTVEYGFVPIGEIVDKGIESSVFSIDSNGIVYTQPIAQWHHRGKQEVFEYSLEDGSIIKATKDHKFMTQDGKMLPIDEIFEQELDLLQVKGLPEGGHHHHHH | 14886 | 14886 |
| HA-H3(1-18)-AVA$^N$ | MYPYDVPDYAGGARTKQTARKSTGGKAPRKCLSYDTEVLTVEYGFVPIGEIVDKGIESSVFSIDSNGIVYTQPIAQWHHRGKQEVFEYSLEDGSIIKATKDHKFMTQDGKMLPIDEIFEQELDLLQVKGLPEGGHHHHHH | 15769 | 15769 |
| HA-H3(1-27)-AVA$^N$ | MYPYDVPDYAGGARTKQTARKSTGGKAPRKQLATKAARKCLSYDTEVLTVEYGFVPIGEIVDKGIESSVFSIDSNGIVYTQPIAQWHHRGKQEVFEYSLEDGSIIKATKDHKFMTQDGKMLPIDEIFEQELDLLQVKGLPEGGHHHHHH | 16737 | 16737 |
| HA-H3(1-31)-AVA$^N$-His | MYPYDVPDYAGGARTKQTARKSTGGKAPRKQLATKAARKSAPACLSYDTEVLTVEYGFVPIGEIVDKGIESSVFSIDSNGIVYTQPIAQWHHRGKQEVFEYSLEDGSIIKATKDHKFMTQDGKMLPIDEIFEQELDLLQVKGLPEGGHHHHHH | 17064 | 17064 |
| His-SUMO-Cys-AVA | MGSSHHHHHHGSGLVPRGSASMSDSEVNQEAKPEVKPEVKPETHINLKVSDGSSEIFFKIKKTTPLRRLMEAFAKRQGKEMDSLRFLYDGIRIQADQTPEDLDMEDNDIIEAHREQIGGCLSYDTEVLTVEYGFVPIGEIVDKGIESSVFSIDSNGIVYTQPIAQWHHRGKQEVFEYSLEDGSIIKATKDHKFMTQDGKMLPIDEIFEQELDLLQVKGLPE | 11579 | 11576 |

Table S3. Primary antibodies used in this manuscript.

| Host | Epitope | WB | IF | *In Vivo* | Vendor |
|---|---|---|---|---|---|
| Chicken | Anti-H3 | 1: 1000 | - | - | Abcam |
| Mouse | Anti-H3 | 1: 1000 | - | - | Abcam |
| Mouse | Anti-HA | 1: 1000 | - | - | Thermo Fisher Scientific |
| Rabbit | Anti-HA | 1: 1000 | - | - | CST |
| Rabbit | Anti-HA | - | - | 1: 500 | Abcam |
| Rat | Anti-HA | - | 1: 125 | - | Roche |
| Mouse | Anti-Flag | - | 1:500 | - | Sigma Aldrich |
| Mouse | Anti-Flag | - | - | 1: 200 | Sigma Aldrich |

Table S4. Secondary antibodies used in this manuscript.

| Host | Epitope | Label | WB | IF | *In Vivo* | Vendor |
|---|---|---|---|---|---|---|
| Donkey | Anti-Chicken | IRDye 800CW | 1: 15000 | - | - | Li-Cor |
| Goat | Anti-Mouse | IRDye 680RD | 1: 15000 | - | - | Li-Cor |
| Goat | Anti-Mouse | IRDye 800CW | 1: 15000 | - | - | Li-Cor |
| Goat | Anti-Mouse | IRDye 568RD | - | 1: 500 | - | Abcam |
| Donkey | Anti- Mouse | Alexa Fluor 647 | 1: 15000 | - | 1:400 | Invitrogen |
| Goat | Anti-Mouse | IRDye 488RD | - | 1: 500 | - | Abcam |
| Goat | Anti-Rabbit | IRDye 800CW | 1: 15000 | - | - | Li-Cor |
| Goat | Anti-Rabbit | IRDye 680RD | 1: 15000 | - | - | Li-Cor |
| Donkey | Anti-Rabbit | Alexa Fluor 488 | 1: 15000 | - | 1:400 | Invitrogen |
| Donkey | Anti-Rabbit | Alexa Fluor 568 | 1: 15000 | - | 1:400 | Invitrogen |
| Goat | Anti-Rat | IRDye 568RD | - | 1: 500 | - | Abcam |
| Goat | Anti-Rat | IRDye 488RD | - | 1: 500 | - | Abcam |
